# Dynamic expression of ASCL1 drives neurogenesis from infant and adult macaque Müller glia into immature retinal ganglion cells

**DOI:** 10.1101/2025.08.21.671552

**Authors:** Marina Pavlou, Juliette Wohlschlegel, Josh Hahn, Lew Kaplan, Fred Rieke, Michael B Manookin, Isabel Ortuño-Lizarán, Marlene Probst, Aric R Prieve, Thomas A Reh

## Abstract

Strategies to stimulate the regeneration of neurons in the adult central nervous system can offer universal solutions for neurodegenerative diseases. Taking lessons from naturally regenerating species, such as the zebrafish, we have previously shown that vector-mediated expression of proneural transcription factors can stimulate neurogenesis from the resident Müller glia (MG) population in the adult mouse retina, both *in vitro* and *in vivo*. To bring this closer to translation, we now show that vector-mediated expression of the proneural transcription factor ASCL1 can reprogram adult macaque MG into functional neurons. To this end, we established purified MG cultures and show they retain a mature transcriptomic profile that correlates with foveal and peripheral MG. Importantly, MG-derived neurons express retinal ganglion cell markers, can fire action potentials and have a transcriptome that overlaps with developing human and adult macaque retinal ganglion cells. To refine this approach for clinical application, we incorporated microRNA-124 target sites in the reprogramming cassette and show that this restricts expression to MG in mixed primary cultures and intact explant cultures of adult macaque retina. Regulating ASCL1 expression with microRNA-124 target sites maintained the reprogramming efficiency from adult MG cultures and improved the yield of RGC-like neurons from infant MG cultures. Most importantly, with this vector cassette we successfully reprogrammed macaque MG from both adult and infant retina into HuC/D+ neurons. Our findings demonstrate that ASCL1 can induce neurogenesis from macaque MG across ages and provide a targeted, effective strategy for potential clinical translation in retinal repair.

## Introduction

Degeneration of neurons in the central nervous system (CNS), namely the brain, retina and spinal cord, leads to permanent deficits in sensory and cognitive function in humans. Although this is the case for humans and other mammals, teleost fish, amphibians, and birds, can activate a regenerative program following neuronal death, which replaces lost neurons and restores function^1–5^. In the retina, this regenerative response is mediated by the upregulation of developmental genes and dedifferentiation of retinal pigmented epithelial (RPE) cells or Müller glia (MG) into proliferating progenitors that then give rise to all neuronal classes^6^. One of the critical factors involved in regeneration in fish is the Achaete-Scute Family BHLH Transcription Factor 1 (ASCL1)^7,8^, and we have shown that ASCL1 overexpression can stimulate neurogenesis in both immature and adult mouse MG ^9–11^. In addition, using both transgenic mouse models and AAV-mediated expression of ASCL1, alone or in combination with other proneural factors, we have obtained functional bipolar, amacrine and RGC-like neurons from adult mouse MG *in vivo*9,^12–14^.

To advance the therapeutic potential of this strategy, we are interested in using proneural factors to reinstate this regenerative capacity in adult non-human primate (NHP) and human MG. Previously we have shown that ASCL1 can stimulate neurogenesis from developing human MG that were isolated from retinal organoids and 3D fetal retina^15,16^. What remains unclear is whether ASCL1 can drive neurogenesis from adult MG in NHPs or humans, and if so, what type of neurons can be regenerated. To address this, we established primary cultures of purified MG from macaque retina across various ages and tested the effects of dynamic ASCL1 overexpression. Adult MG cultures were successfully reprogrammed into neurogenic progenitors and retinal neurons after overexpressing ASCL1 using a strong ubiquitous promoter. Macaque glia-derived neurons express retinal ganglion cell (RGC) proteins, share electrophysiological properties with immature RGCs and have a similar transcriptome to developing human and adult macaque RGCs.

To tailor this strategy for translation, we incorporated miR124 target sites at the 3’UTR of the reprogramming cassette in order to: (1) enhance the specificity of the reprogramming cassette by de-targeting endogenous neurons and (2) test whether dynamically regulating ASCL1 expression affects neurogenesis from MG. We show that this strategy effectively restricts gene expression to MG in both dissociated cultures and intact retinal explants. Importantly, vectors expressing ASCL1 with or without a miRNA de-targeting element yield the same reprogramming efficiency from adult MG. Interestingly, however, the dynamic regulation of ASCL1 significantly improved the yield of RGC-like neurons from infant MG. Taken together, our results support further clinical translation of this approach as a potential regenerative medicine for retinal repair.

## Results

### Short-term cultures of adult primate Müller glia retain mature profile

To determine whether adult macaque (*Macaca nemestrina*) MG can be reprogrammed with proneural TFs, we used dissociated cultures of retina. We previously established *in vitro* models of Müller glia culture from various sources, such as (1) postnatal day 12 mouse retina^10^ and (2) human organoid cultures and human fetal retina^15^. These culturing strategies relied on consecutive passaging of primary retina to eliminate surviving neurons and obtain a glia-rich culture for subsequent experiments. This approach is possible when the cell source is immature retina, as cells still retain their proliferative capacity *in vitro* (for detailed characterization of P12 mouse cultures see^17^; however, establishing MG cultures from adult retina is more challenging since adult cells have a very limited proliferation capacity, as observed in mice^18^ and macaques.

We initially used a similar dissociation approach, as previously described, to culture MG from adult macaque retina that was divided into quadrants to assess any regional differences. Adult primate eyecups were collected within an hour postmortem and the retina was isolated from the rest of the tissue for subsequent dissociation. After dissociation, the cells were plated in Matrigel-coated wells and cultured to confluence. We found that MG from all regions of the retina could be expanded using this method (Suppl Fig 1A-C). The majority of the cells could be labeled with antibodies for MG specific proteins, like CRALBP, thereby confirming that we could culture MG irrespective of the retinal region, in instances where we could not obtain the entire globe.

Nevertheless, these primary cultures were soon dominated by other cell types that are highly proliferative, such as SOX2+GFAP+CRALBP-astrocytes (Suppl Fig 1C). As such, we sought to selectively enrich for MG prior to plating so as to eliminate contaminating cell types and limit their culture period.

To this end, we implemented a strategy to enrich for primary MG by adapting a previously established method based on magnetic activated cell sorting (MACS)^19^. Adult macaque retina was dissected and enzymatically dissociated, after which MG were positively selected using CD29 antibody-coated beads (Fig 1A). Due to the gentle nature of this protocol, MG were able to survive, retain their morphology and effectively expand in culture (Fig 1B). This allowed us to enrich for MG and eliminate surviving neurons from the cultures, which were mostly OTX2+ bipolar cells/photoreceptors and sparse HuC/D+ amacrine/retinal ganglion cells (RGCs) (Fig 1C). To determine the degree of purification from the MACS sorting, we used immunolabeling of the CD29+ and CD29-fractions immediately post-MACS (Figure 1D, Suppl Fig 2A-B) and after 4 days *in vitro* (DIV) (Figure 1E-F, Suppl Fig 2C), as well as transcriptomic analysis (Fig 1G-H). Cells in the CD29+ fraction expressed MG markers CRALBP, GS and SOX9 immediately post-MACS (Fig 1D), with some also expressing

**Figure 1:**
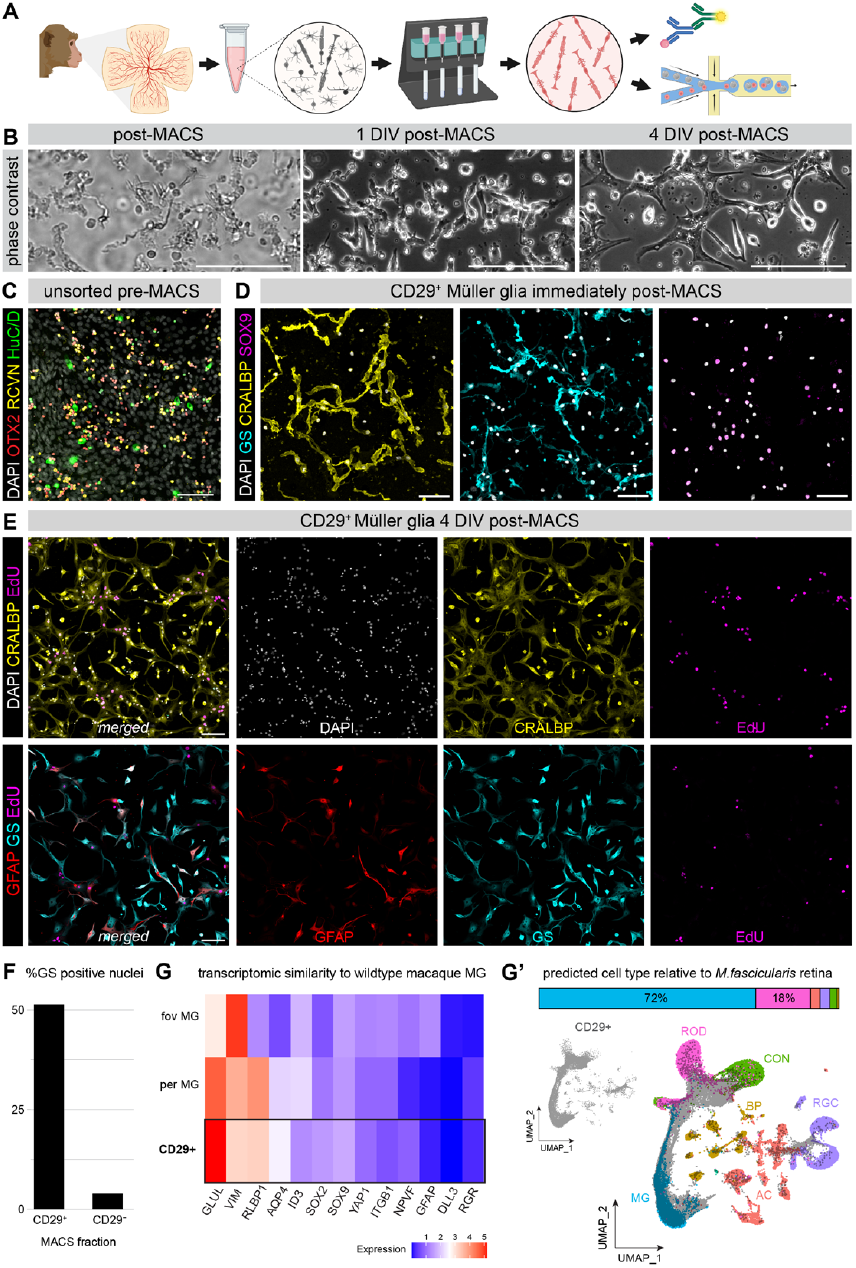
Purified MG cultures from adult macaque retina: (A) schematic diagram of the experimental pipeline used to assess culture composition after magnetic-activated cell sorting adult macaque retina, (B) phase contrast images of the CD29+ fraction over time showing from left to right cells immediately post MACS, 1 day *in vitro* (DIV) and 4 DIV, (C) fluorescence image of an unsorted primary retinal culture showing surviving neurons, (D) fluorescence images of the CD29+ fraction labeled with MG markers immediately post MACS, (E) fluorescence images of the CD29+ fraction labeled with MG markers and EdU after 4 DIV, (F) bar plot showing the percentage of GS+ cells in the CD29+ and CD29-fractions post MACS, (G) heatmap of gene expression levels for MG markers using single cell RNA-sequencing (scRNA-seq) data to compare the CD29+ fraction of *Macaca nemestrina* to the *Macaca fascicularis* peripheral and foveal MG published in ^20^, (G’) alluvium plot showing the distribution of predicted cell type labels from the *Macaca fascicularis* whole retina scRNA-seq onto the CD29+ fraction of *Macaca nemestrina* and UMAPs of the integrated data showing the CD29+ cells alone on the left and the two overlapping samples on the right with cell clusters labelled as per ^20^. Scalebars: 200μm for (B), 50μm for (C-D), 100μm for (E).

GFAP (Suppl Fig 2B). After 4 DIV a subset of cells proliferated (EdU+) while still retaining the same MG markers CRALBP, GS, GFAP (Fig 1E) and CD29 (Suppl Fig 2C). MACS eliminated most of surviving neurons as there were no detectable BRN3+ or HuC/D+ RGCs in the CD29+ fraction of MG, though a small number of OTX2+ neurons persisted (Suppl Fig2B). By harvesting MG directly from the macaque retina and culturing them for a very short time (4 DIV), we were able to obtain a MG population that transcriptionally resembles adult MG from both the fovea and periphery of previously published *Macaca fascicularis* retina^20^(Fig 1G). When we mapped the CD29+ fraction of *Macaca nemestrina* cells to the *Macaca fascicularis* reference atlas^20^, the cells largely overlapped with the MG cluster (Fig 1G’, UMAPs). We also used label transfer to assign cell type labels from the reference atlas to the CD29+ sorted cells. Of the cells that were confidently assigned to a class label, a majority were assigned to be MG, with the primary contaminating cells being rod photoreceptors (∼ 10%) (Fig 1G’, bar graph). This culture system of MG was therefore an appropriate testbed for subsequent reprogramming experiments.

### Adult Müller glia are reprogrammed into retinal ganglion-like cells with ASCL1

To test the responses of adult MG to ASCL1 overexpression, MACS-purified MG cultures were transduced with viral vectors expressing ASCL1-eGFP or eGFP alone under the control of a ubiquitous CMV promoter (Fig 2A). Within two days post infection (DPI) there was widespread reporter expression (Fig 2B, Suppl Fig 3A) and by 7 DPI there were evident morphological changes to the eGFP+ cells between the two treatments (Fig 2B). Single-cell RNA sequencing (scRNA-seq) at 7 DPI confirmed a significant change in the transcriptome of transduced cells, with new clusters generated following ASCL1 overexpression compared to control (Fig 2C-C’). UMAP plots of the integrated data from both ASCL1-eGFP and eGFP treatments show several distinct clusters that we assign as MG, cycling cells, Neurogenic Precursors (NPre), and MG-derived retinal ganglion cells (MG→RGCs), based on known marker genes for these cell types (Figure 2E). The cycling cells, NPre and MG→RGCs are only present in the ASCL1 condition, consistent with neural reprogramming. In addition, we saw small clusters of rod photoreceptors and bipolar cells present in both treatment groups and presume these are carried over during MACS as described above.

**Figure 2:**
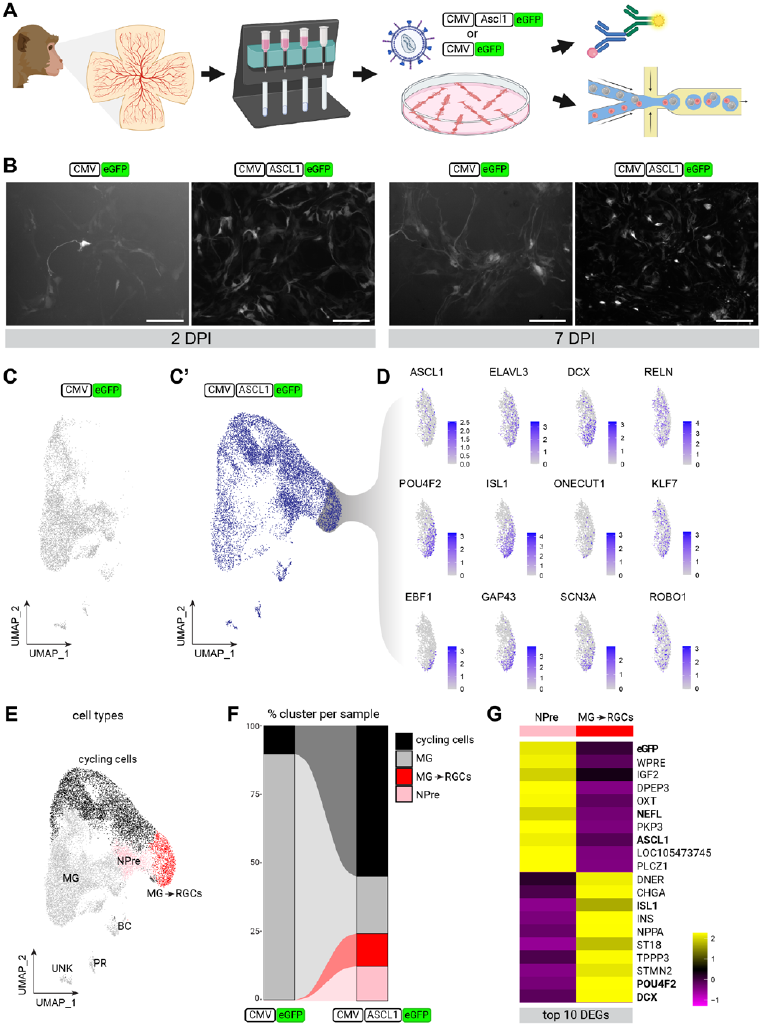
ASCL1-mediated reprogramming of adult macaque MG: (A) schematic diagram of the experimental pipeline used to assess the effects of ASCL1 overexpression on adult MG, (B) fluorescence images of live MACS-purified MG cultures at 2 days post infection (DPI) and 7 DPI treated with lentiviruses expressing either CMV-eGFP or CMV-ASCL1-eGFP, (C-C’) UMAPs of integrated scRNA-seq data from 7DPI cultures split by treatment as annotated above each UMAP, (D) feature plots showing gene expression levels for retinal ganglion cell (RGC) markers in unique cluster generated after MG treatment with CMV-ASCL1-eGFP, (E) UMAP of integrated scRNA-seq data from 7DPI cultures showing cell clusters identified based on common transcripts, (F) alluvium plot showing the percentage of clusters: cycling cells, MG, neurogenic precursors (NPre) and MG-derived RGCs (MG→RGCs) per sample, (G) heatmap of top 10 differentially expressed genes (DEGs) between clusters NPre and MG→RGCs with selected genes of interest in bold. Scalebars: 200μm for (B).

The three main clusters induced by ASCL1 show strong evidence of the stages in reprogramming previously shown in mouse retina: (1) ASCL1 stimulates MG proliferation and cells with markers of mitotic proliferation (cycling cells); (2) ASCL1 increases downstream targets HES6 and DLL1 (NPre) while RLBP1 is downregulated (Suppl Fig 3B); (3) ASCL1 induces MG to generate cells with the transcriptome of neurons (MG→RGCs). The NPre and MG→RGCs clusters were exclusively formed after CMV-ASCL1-eGFP treatment compared to CMV-eGFP control (Fig 2F), with the latter expressing neither ASCL1 transcripts nor protein (Suppl Fig 3C).

Interrogating the neuron cluster more closely, we observed transcripts for several RGC markers, such as POU4F2, ELAVL3, DCX, ISL1, ONECUT1, KLF7, EBF1, GAP43, SCN3A,RELN and ROBO1 (Fig 2D). Interestingly, we saw the upregulation of RGR, a nonvisual opsin gene expressed in foveal MG ^21,22^, cone photoreceptors and a subset of RGCs^23^(Suppl Fig 3B). An unbiased query of the top differentially expressed genes between NPre and MG→RGCs confirmed the distinct expression of genes ISL1, POU4F2 and DCX (Fig 2G), verifying that when ASCL1 is expressed in adult macaque MG it stimulates the neurogenesis of RGC-like neurons.

As noted above, there were small clusters of photoreceptors and bipolar cells detected for both treatments in the scRNA-seq data, which were carried over during MACS (Fig 2E); this is a phenomenon we have encountered and addressed in previous studies where we used fluorescence-activated cell sorting (FACS) to isolate lineage-traced glia^12,13,16,24^. To confirm that the RGC-like neurons are glia-derived, we used EdU incorporation and immunocytochemistry. Immunolabeling of transduced cells corroborated the scRNA-seq outcome, with eGFP+ cells expressing ASCL1, ISL1 (Suppl Fig 3D), BRN3 and HuC/D (Fig 3A). Importantly, a subset of these cells were also EdU+ (white arrows in Fig 3A, Suppl Fig 3D), confirming that these are *bona fide* newborn neurons.

**Figure 3:**
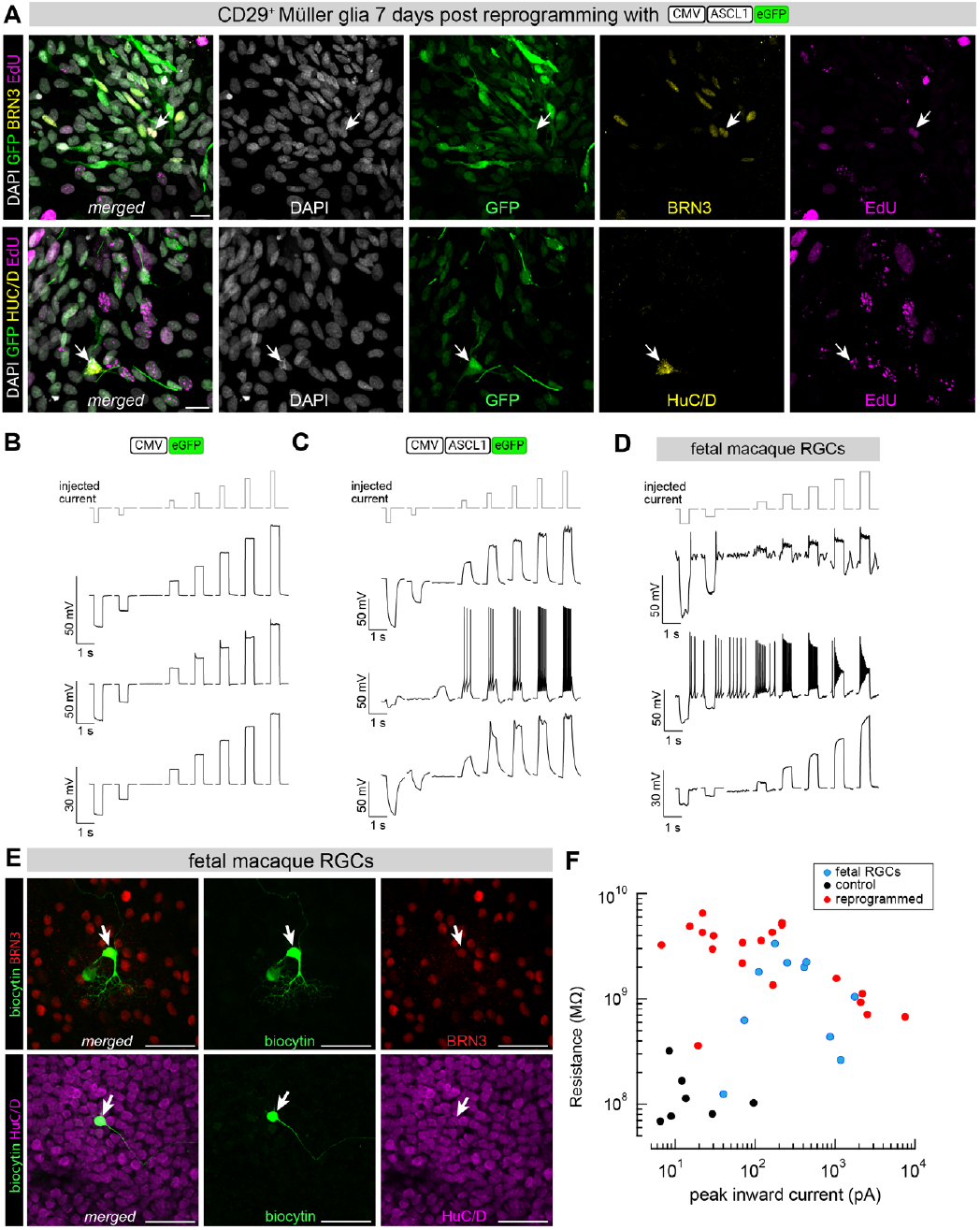
ASCL1 reprograms adult macaque MG into functional immature RGCs: (A) fluorescence images of MACS-purified MG cultures treated with CMV-ASCL1-eGFP and fixed at 7DPI with white arrows showing examples of GFP+EdU+ cells that co-label with RGC markers BRN3 (top panel) or HuC/D (bottom panel), (B-C) patch-clamp recordings of voltage responses to incremental current steps for three representative cells treated with (B) CMV-eGFP or (C) CMV-ASCL1-eGFP, (D) patch-clamp recordings of voltage responses to incremental current steps for three representative fetal macaque RGCs, (E) fluorescence images of whole-mount fetal macaque retina showing examples of patch-clamped and biocytin-filled RGCs that are either BRN3+ (top panel) or HuC/D+ (bottom panel), (F) scatterplot summarizes the peak inward current to membrane resistance relationship for patch-clamped adult MG after 7DPI (n=7 black circles: control CMV-eGFP, n=20 red circles: reprogrammed CMV-ASCL1-eGFP) and endogenous fetal RGCs from day 155 retina (n=10 blue circles). Scalebars: 20μm for (A), 50μm for (E).

To profile the electrophysiological properties of these reprogrammed neurons, we performed patch-clamp recordings of GFP+ cells that were transduced either with the control CMV-eGFP vector (n=7 cells) or with CMV-ASCL1-eGPF (n=20 cells) at 7 DPI. Cells transduced with CMV-eGFP showed no spiking in response to incremental current steps (three examples, Fig 3B) and had electrical properties resembling native Müller glia, including low input resistance and a lack of inward current elicited by a voltage step. Cells transduced with CMV-ASCL1-eGFP (example dye-filled cells, Suppl Fig 3E) varied considerably but consistently showed higher input resistance than the control cells. Some of the reprogrammed cells fired repetitive action potentials (three examples, Fig 3C) and many had significant inward currents in response to a voltage step. To correlate these responses to endogenous RGCs, we also performed patch-clamp recordings of young RGCs from day 155 fetal macaque retina (three examples, Fig 3D). Similar to the reprogrammed RGCs, endogenous fetal RGCs had variable responses, with some firing action potentials with peak amplitudes of +10 or +20 mV and some failing to generate action potentials. To visualize the cells, fetal RGCs were filled with biocytin, and the tissue was subsequently labelled for markers BRN3 and HuC/D. We obtained two examples of BRN3+ and HuC/D+ fetal RGCs (white arrows, Fig 3E) whose morphology suggests they are likely midget cells. Some of the recorded cells were BRN3-with branched dendritic arbors that resemble parasol RGCs (Suppl Fig 3F). Membrane resistances and inward currents for all recorded cells were summarized, showing an increased resistance of reprogrammed cells (n=20 cells, red circles, Fig 3F) compared to control (n=7 cells, black circles, Fig 3F) and a similar distribution of responses to fetal RGCs (n=10 cells, blue circles, Fig 3F).

Although the morphology of MG-derived RGCS after 7 days doesn’t resemble mature RGCs, if cells are left in culture for longer i.e. 21 days, they begin to acquire more complex neuronal morphologies (Fig 4A). This development in RGC morphology is accentuated when waiting even longer, with examples of glia-derived neurons (GFP+BRN3+TUBB3+) surviving up to 5 months post infection (Fig 4B). Since extending the survival time of the glial-derived RGCs led to more complex morphologies, we wondered whether longer survival times would also lead to maturation of their transcriptome (Fig 4C). To address the extent and direction of maturation, we plotted the transcriptome of reprogrammed macaque MG from both 7 DPI and 21 DPI against the developing human retina^15^ (fetal weeks 8 and 10) and used label transfer to assign the reprogrammed macaque MG an equivalent developmental stage (Fig 4D-D’’). The macaque MG and cycling cells clustered proximally with the corresponding cell types in the human fetal retina, i.e. MG and cycling multipotent progenitors (MPC). ASCL1+ NPre from macaque cultures clustered proximally to human MPC, NPre and neuronal branches (Fig 4D’). Finally, macaque MG→RGCs displayed signs of neuronal identity, with 47% of cells being assigned a neuronal precursor label and 39% clustering with the branches of developing RGCs, amacrine and horizontal cells, likely due to shared genes amongst these branches (Fig 4D’’). If we compare our *Macaca nemestrina* ASCL1+ NPre and MG→RGCs scRNA-seq data (black outline, Fig 4E) to the published transcriptomic profile of RGC subtypes in the *Macaca fascicularis* retina^20^, there are common upregulated genes such as GRIA4, SEMA6A, EYA1, ISL1 and SPP1 (Fig 4E); however, no single subtype of RGCs was generated by the ASCL1-expressing MG.

**Figure 4:**
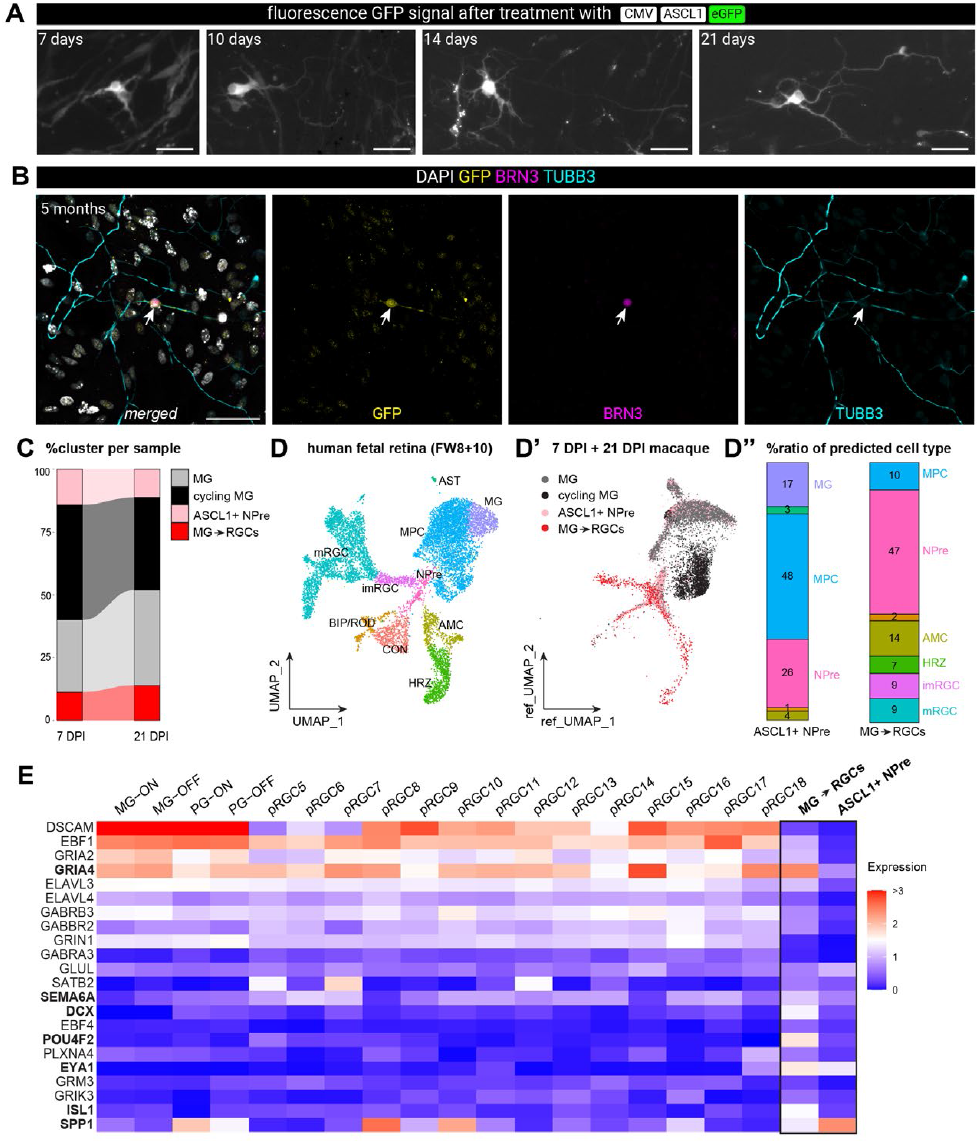
Long-term cultures of MG-derived RGCs: (A) fluorescence images of live MACS-purified MG cultures at 7-, 10-, 14- and 21-days post infection (DPI) with LV/CMV-ASCL1-eGFP showing changes in the morphology of GFP+ cells, (B) fluorescence images of fixed MACS-purified MG culture at 5 months post infection with LV/CMV-ASCL1-eGFP with arrow indicating a reprogrammed GFP+BRN3+TUBB3+ cell, (C) alluvium plot showing the percentage of clusters: MG, cycling MG, ASCL1+ neurogenic precursors (NPre) and MG-derived RGCs (MG→RGCs) per sample either 7 DPI or 21 DPI, (D-D’’) transcriptomic comparison of reprogrammed cultures of adult macaque MG to human fetal retina, (D) UMAP of merged human fetal retina from gestation week 8 and 10 clustered by cell type as in ^15^, (D’) UMAP of macaque MG, cycling MG, ASCL1+ NPre and MG→RGCs distributed based on their transcriptomic similarity to the reference coordinates of the human fetal retina in (D), (D’’) alluvium plots showing the percentage ratio of predicted cell types following label transfer from human fetal retina onto the ASCL1+NPre (left bar) and MG→RGCs (right bar), (E) heatmap of gene expression for the most common RGC markers across all macaque RGC subtypes as identified in ^20^ and our reprogrammed macaque cells highlighted with a bold margin. Scalebars: 50μm

### A miRNA-mediated de-targeting strategy facilitates MG reprogramming in intact retina

Having verified the potential for ASCL1 overexpression to drive neurogenesis from adult macaque MG, we proceeded to tailor this approach for *in vivo* application. For that, we needed a strategy that would restrict vector-mediated ASCL1 expression in MG. This is a particular challenge since even in smaller mammals such as mice, an all-in-one vector that expresses proneural factors under a glia-specific promoter can lead to ectopic expression in endogenous neurons^25^. As such, we opted for harnessing the natural silencing capacity of miRNAs to de-target the vector cassette from endogenous neurons and drive its expression only in glia. To this end, we queried Multiome data of the developing human retina and picked out the well-known miR-124, which is expressed in all newborn neurons but not MG (Suppl Fig 4A-A’’).

We generated two new lentiviral vector casettes with the ubiquitous CMV promoter driving gene expression and inserted miR-124 target sites at the 3’UTR. To assess neuronal de-targeting, primary retinal cultures from macaque retina were transduced with the CMV-mNG-miR124T reporter vector. These cultures were obtained from dissociated retina that did not undergo MACS and were allowed to attach for 4 DIV before transduction with the CMV-mNG-miR124T reporter vector. Cultures were incubated for 7 days, which is the same timeline for all previous reprogramming experiments, and then fixed for ICC. We found no overlap between TUBB3+ neurons and the vector reporter mNG, indicating that the de-targeting strategy was effective (Suppl Fig 4B-C). Furthermore, we tested whether the addition of miR-124 target sites would interfere with the conversion of glia into neurons by transducing MACS-purified MG with a vector carrying a CMV-ASCL1-mNG-miR124T or a CMV-mNG-miR124T control cassette (Fig 5A). After 7 DPI, MG transduced with CMV-ASCL1-mNG-miR124T expressed ASCL1 and the RGC marker BRN3, whereas those transduced with the control vector did not (Fig 5B). The addition of miR-124 target sites did not affect the number of ASCL1+ cells (4.28% versus 4.29% of all sequenced cells) or neurogenesis yield from adult macaque MG (Fig 5C-D). All cell clusters previously identified after 7 DPI were present at equal frequency after reprogramming with the CMV-ASCL1-mNG-miR124T vector (Fig 5D’). Furthermore, when performing pseudotime analysis of glia-to-neuron reprogramming that is superimposed on the developing human fetal retina (Fig 5E), the cell density across pseudotime was the same for both vectors expressing ASCL1 with or without miR-124 target sites (Fig 5F). This confirms that a miR124-mediated de-targeting strategy does not interfere with glia-to-neuron reprogramming, which is an essential prerequisite for applying this strategy on intact macaque retina.

**Figure 5:**
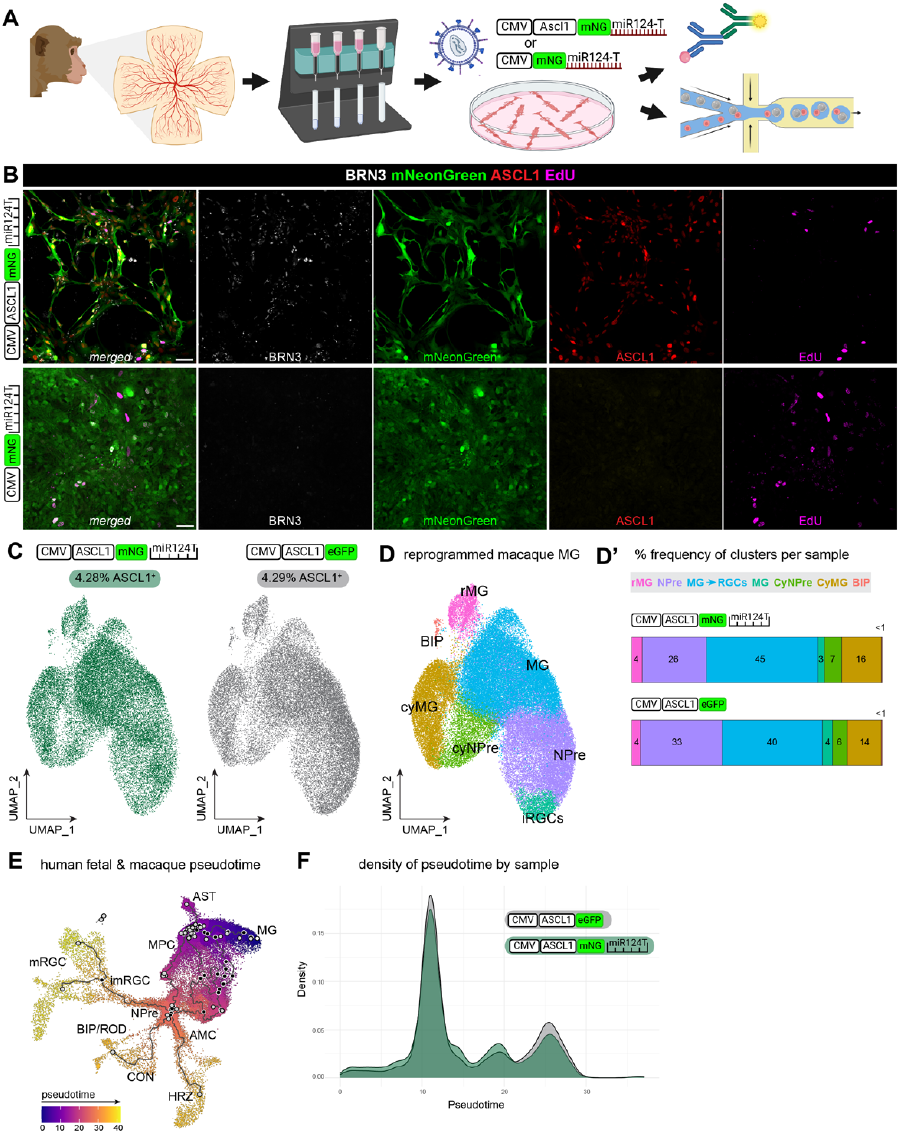
Addition of miR124 target sites does not affect the reprogramming of adult macaque MG: (A) schematic diagram of the experimental pipeline used to assess the effects of ASCL1 overexpression on adult MG with new constructs that include miR124 target sites at the 3’UTR, (B) fluorescence images of MACS-purified MG cultures treated with CMV-ASCL1-mNG-miR124T (top panel) or control CMV-mNG-miR124T (bottom panel) and fixed at 7DPI, (C) UMAPs of integrated scRNA-seq data from 7DPI cultures treated with either CMV-ASCL1-mNG-miR124T or CMV-ASCL1-eGFP split by treatment as annotated above each UMAP with the percentage of ASCL1+ cells for each dataset noted above the UMAP, (D) UMAP of integrated scRNA-seq data from 7DPI cultures showing cell clusters identified based on common transcripts (D’) alluvium plots showing the percentage of each cell cluster in each dataset, (E) UMAP of integrated scRNA-seq data from the human fetal retina and reprogrammed adult macaque samples showing pseudotime nodes from earliest to latest cell types, (F) distribution of cell density from reprogrammed adult macaque samples across pseudo-time.

To further confirm the glial specificity and test the reprogramming potential of the miRNA de-targeting vectors, we used explant cultures from intact adult macaque retina, prepared as previously described ^26,27^. On the day of culture, the explants were transduced with LV vectors CMV-ASCL1-mNG-miR124T or CMV-mNG-miR124T. After three weeks of incubation, we observed widespread labeling of mNG+ cells for both vectors. Explants treated with CMV-ASCL1-mNG-miR124T had robust ASCL1 expression in mNG+ cells that were mostly SOX2+ glia (Fig 6A). ASCL1 expression induced a subset of glia to re-enter the cell cycle or generate neurons, i.e. co-label with EdU or AC/RGC marker HuC/D, respectively(white arrows, Fig 6A). Importantly, neither glial proliferation nor reprogramming was observed in explants treated with the control vector CMV-mNG-miR124T, which elicited mNG expression only in SOX2+ glia (Fig 6B). Of a total 300 mNG+ cells counted from explants treated with CMV-ASCL1-mNG-miR124T, 90.5% were SOX2+ and of those, 20.6% were SOX2+EdU+ (left panel, Fig 6C). Importantly, 100% of mNG+ cells were ASCL1+ and of those, 5.3% were ASCL1+EdU+ and 5.4% were ASCL1+HuC/D+ (right panel, Fig 6C). In contrast, from the total 79 mNG+ cells counted from explants treated with CMV-mNG-miR124T, 100% were SOX2+ and there were no mNG+ cells found that expressed ASCL1, EdU or HuC/D (Fig 6D).

**Figure 6:**
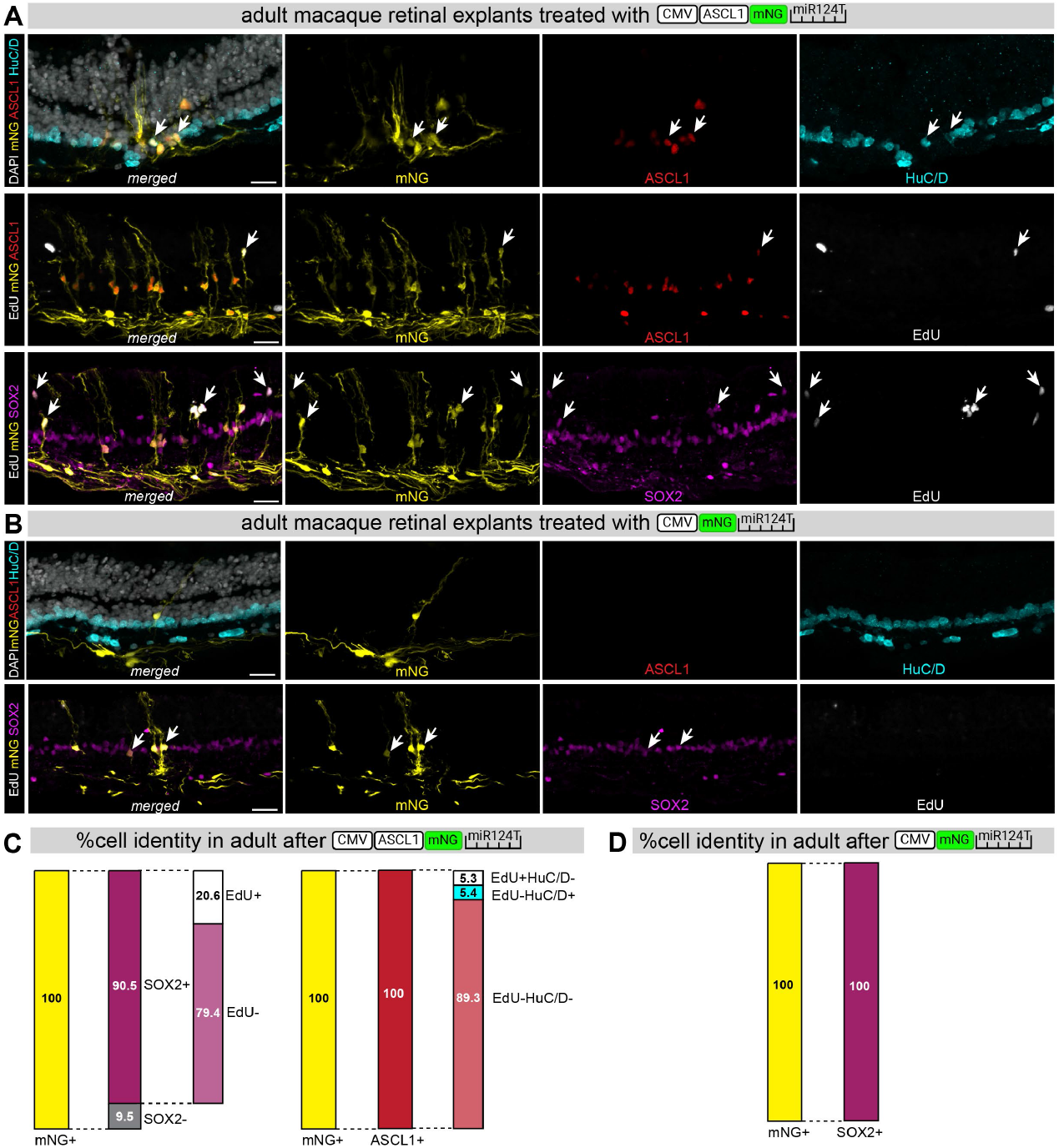
MG-derived neurogenesis in adult macaque retinal explants: (A) fluorescence images of tissue sections from macaque retinal explants treated with lentiviral vector expressing CMV-ASCL1-mNG-miR124T for 3 weeks *in vitro* with white arrows showing examples of mNG+ASCL1+HuC/D+ cells (top panel), mNG+ASCL1+EdU+ cells (middle panel), and mNG+SOX2+EdU+ cells (bottom panel), (B) same as (A) but for retinal explants treated with control lentiviral vector CMV-mNG-miR124T showing the absence of ASCL1 and HuC/D labelling in mNG+ cells (top panel) and white arrows showing examples of mNG+SOX2+EdU-cells (bottom panel), (C) staggered bar plots showing the percentage of cell identity quantified from fluorescent images of retinal sections after treatment with CMV-ASCL1-mNG-miR124T, (D) same as (C) but for retinal explants treated with control lentiviral vector CMV-mNG-miR124T. Scalebars: 20μm

### Dynamic regulation of ASCL1 via miRNA de-targeting improves neurogenesis yield in an age-dependent manner

The results presented so far show that ASCL1 can stimulate neurogenesis from MG in adult macaque retina from animals aged between 3-years and 2-months old to 22-years old (Table 1). In mice, MG from younger mice have a greater neurogenic potential than older animals ^9,11,28^ and so we wondered whether this phenomenon would also occur in non-human primates. To this end, infant macaque retina was harvested from a 7-month-old animal that failed to thrive and was processed similarly to the adult to establish MG cultures (Fig 7A). The CD29+ fraction following MACS was enriched for GS+ MG, though a carryover of RCVRN+ cells was also detected, as in the adult (see above) (Fig 7B). Enriched MG cultures were then transduced with the vectors used previously in the adult macaque and processed for scRNA-seq after 7 DPI (Fig 7C). All four datasets were integrated, and clusters were called based on common gene expression profiles, namely reactive MG, MG, cycling MG, NPre, cycling NPre, and MG→RGCs. The cycling NPre and MG→RGCs clusters were exclusive to the samples treated with ASCL1-expressing vectors, whereas control samples had a higher proportion of GFAP+ reactive MG (Fig 7C-D). Strikingly, infant MG transduced with the CMV-ASCL1-mNG-miR124T cassette had a 3-fold greater yield of glia-derived RGCs and a 4-fold lower proportion of cycling NPre compared to the CMV-ASCL1-eGFP cassette (MG→RGCs: 10% versus 3%; cyNPre: 4% versus 16% of total cells) (Fig 7D). This suggests that the presence of miR-124 target sites likely facilitated a faster cell-cycle exit for cells to become immature RGCs.

**Table 1:** summary of non-human primate (NHP) samples (species: *Macaca nemestrina*) and treatments; MACS: magnetic activated cell sorting, LV: lentiviral transduction, ephys: path-clamp recordings, SCR: single cell RNA-sequencing, TCR: targeted cell recovery.

| ID | Age (y,m,w,d) | Treatment | SCR (y/n) | SCR ID | TCR |
| --- | --- | --- | --- | --- | --- |
| NHP50 | 22y | MACS + LV | Y | SCR 198, 199 | 8,000 |
| NHP51 | 4y | MACS + LV | N |  | N/A |
| NHP52 | 10y 7m | MACS + LV | N |  | N/A |
| NHP55 | 5y | MACS + LV + ephys | N |  | N/A |
| NHP57 | 5y 5m | MACS + LV | Y | SCR 226, 231-2 | 8,000 |
| NHP58 | 9y | MACS + LV | N |  | N/A |
| NHP59 | 3y 2m | MACS + LV | N |  | N/A |
| NHP61 | 18y6m | Explant+LV | N |  | N/A |
| NHP62 | 6y 2m | MACS + LV | Y | SCR 241, 242 | 8,000 |
| NHP63 | 7w | MACS + LV | Y | SCR 246-9 | 8,000 |
| NHP65 | 7y 11m | Explant+LV | N |  | N/A |
| NHP66 | 155d | ephys or explant+ LV | N |  | N/A |

**Figure 7:**
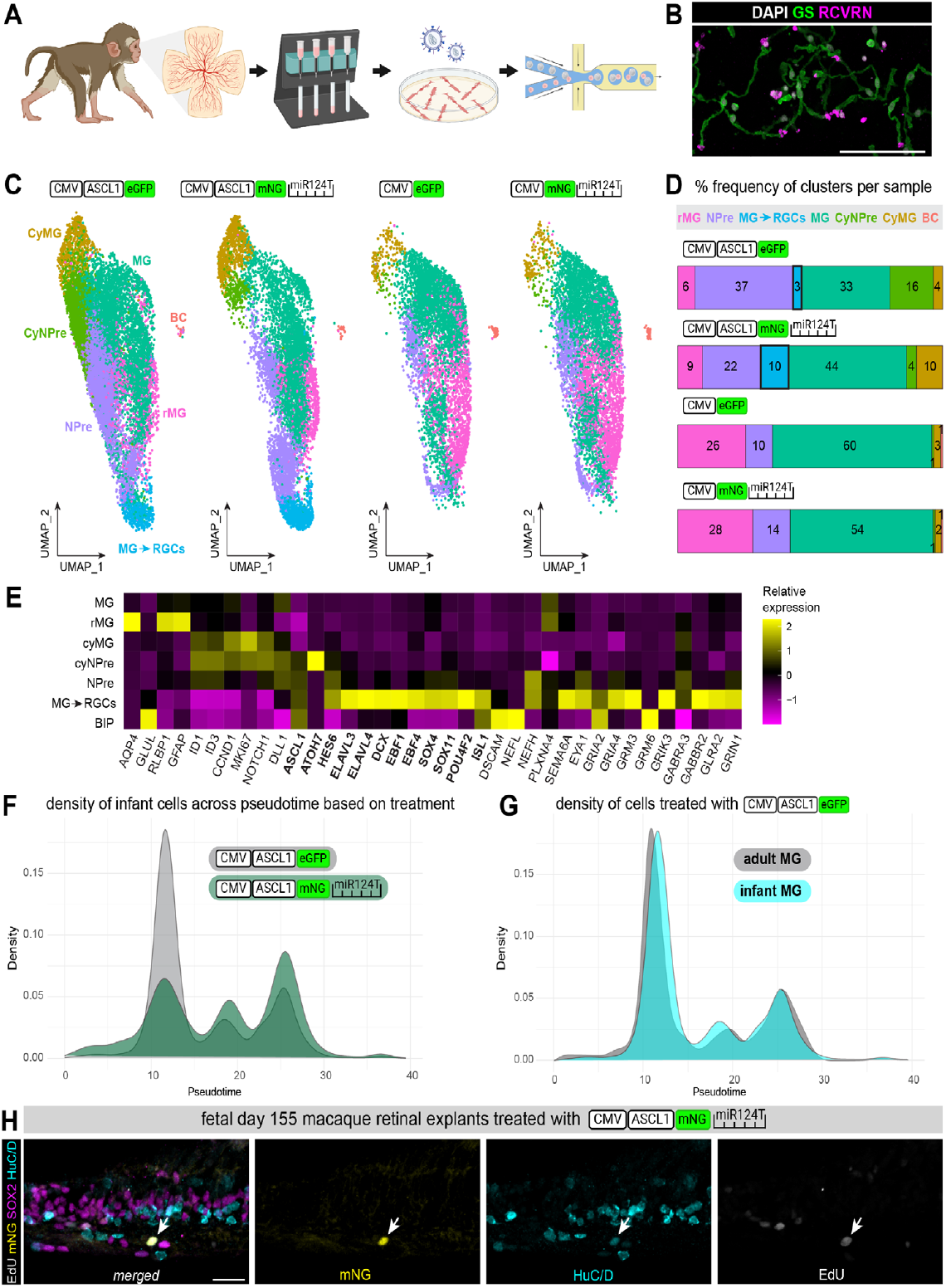
Infant macaque MG are reprogrammed into immature RGCs with ASCL1: (A) schematic diagram of the experimental pipeline used to assess the effects of ASCL1 overexpression on infant MG, fluorescence images of the CD29+ fraction labeled with MG marker GS and bipolar/photoreceptor marker RCVRN immediately post MACS, (C) UMAPs of integrated scRNA-seq data from 7DPI cultures split by treatment as annotated above each UMAP and with cell clusters identified based on common transcripts, (D) alluvium plots showing the percentage of each cell cluster in each dataset with MG→RGCs highlighted with a bold margin, (E) heatmap of gene expression for relevant markers across all cell clusters identified in (C), (F) distribution of cell density from reprogrammed infant macaque samples treated with either CMV-ASCL1-eGFP or CMV-ASCL1-mNG-miR124T across pseudotime, (G) distribution of cell density from either adult or infant macaque samples treated with CMV-ASCL1-eGFP, (H) fluorescence images of tissue sections from macaque retinal explants treated with lentiviral vector expressing CMV-ASCL1-mNG-miR124T for 3 weeks *in vitro* with white arrows showing example of mNG+HuC/D+EdU+ cell. Scalebars: 100μm for (B), 20μm for (H).

This increased neurogenesis was reflected in the relative levels of gene expression across clusters (Fig 7E), where cycling MG and cycling NPre follow the same pattern except the latter expressed higher levels of ASCL1 and ATOH7. Infant MG-derived RGCs expressed several neuronal markers of the RGC fate, as in the adult macaque, including ELAVL3/4, EBF1, EBF4, POU4F2, ISL1, SEMA6A, EYA1 and sodium channel subunits (Fig 7E). Most importantly, the cell density across pseudotime showed a faster transition of cells from glia-to-neurons when ASCL1 was dynamically regulated with miR-124 target sites (Fig 7F). This was not due to the more plastic nature of infant MG relative to adult MG, as MG from both infant (7-weeks old) or adult retina (6 years 2 months old) had the same pseudotime density plot following CMV-ASCL1-eGFP expression (Fig 7G). A slightly younger retina from a fetal day 155 macaque (gestation is at day 183) was used to assess the effect of CMV-ASCL1-mNG-miR124T and after 3 weeks incubation we detected newborn mNG+EdU+HuC/D+ neurons (white arrow, Fig 7H). This was the only explant sample where we detected EdU+HuC/D+ neurons, which we attribute to the accelerated transition from precursors to neurons. Overall, we conclude that overexpressing ASCL1 in macaque MG, whether infant or adult, and dynamically regulating its expression with a miR124T, will result in neurogenesis of immature RGCs.

## Discussion

Reinstating the regenerative capacity of adult MG can have tremendous impact on how patients with retina degeneration are treated. We have learnt a lot from studying how other species respond to regenerative cues but identifying effective strategies on adult tissue that is evolutionarily closest to humans is essential for successful translation. Of the animal models available for research, non-human primates share the most genetic and physiological features with humans (common superfamily: catarrhine primates).

We show that MG from adult macaques can be reprogrammed by the proneural transcription factor ASCL1 to generate progenitor-like cells that produce neurons. This is possible across a wide range of ages, from infant (<12 months), juvenile (1-4 years), adult (4-15 years) and geriatric macaques (>15 years)^29^, as the macaque retina we harvested was from animals aged between 7 weeks and 22 years old (Table 1). MG-derived neurogenesis did not require an HDAC inhibitor in 2D cultures of enriched MG. However, explant cultures were treated with the HDAC inhibitor sodium butyrate which we identified was necessary in previous experiments^9^. Previous studies have shown that vector-mediated ASCL1 expression was insufficient to convert mouse embryonic fibroblasts^30^ or MG from 3D human retina^16^ into RGCs. This could be attributed to the cell species, experimental design, as well as promoter strength, since previous work used either a tetracycline transactivator element or a cell-type specific promoter to drive ASCL1.

Given that the supply of intact macaque retina can be rare and unpredictable, we sought to establish a method for harvesting adult MG and culture them very briefly before testing the effects of ASCL1 overexpression. Several methods have been implemented to enrich for MG from either wildtype or transgenic sources; for an extensive review see^31^. *In vitro* cultures of primary macaque retina has helped make several advancements in understanding physiology and function to an otherwise inaccessible tissue. Initial studies using *in vitro* primary cultures (*Macaca fascicularis* and *Papio cynocephalus*) were able to keep cells viable for a few hours, long enough to perform whole-cell patch-clamp recordings and characterize cell morphology by phase-contrast microscopy^32^. Some reports have claimed to achieve long-term cultures of primary MG from primate origins (*Macaca fascicularis*) ^33,34^ but these have questionable relevance to *in vivo* MG since proteomic analysis of primary MG cultures has demonstrated the dedifferentiation of MG to a fibroblast-like phenotype in culture after ∼3 days *in vitro*^35^. In our experience, primary MG cultures from macaque retina (*Macaca nemestrina*) have a very limited proliferation window that does not extend past one passage. As such, we adapted a published protocol for isolating MG from primary human retina^19^ and kept them in culture for only a limited period, namely 4 days, before transduction with reprogramming vectors. These MG cultures were depleted of neurons, expressed mature MG markers and retained a mature transcriptomic profile that overlapped with adult macaque MG from intact retina.

We used a ubiquitous promoter to drive the expression of ASCL1 in purified cultures of MG and obtained RGC-like neurons within 7 days of incubation. These neurons express RGC markers and some originated from ASCL1+ MG that de-differentiated into cycling progenitors before becoming RGCs. MG transduced with CMV-ASCL1-eGFP had a distinct electrophysiological profile that was characteristic of RGCs and differed significantly from control cells lacking ASCL1. Some even fired action potentials which is a unique feature of RGCs in the retina. Compared to fetal RGCs from a 155-day old macaque retina (gestation 183 days), reprogrammed RGCs had a similar membrane resistance to increments of injected current and expressed the same RGC markers i.e. BRN3 and HuC/D. When mapped onto the developing human fetal retina, we see an overlap with mostly neurogenic precursors and the neuronal branches, particularly of the amacrine and RGC lineages. This matches the fact that no specific RGC subtypes are generated with ASCL1 expression alone during development^36^ and is consistent with the observation that long term cultures show increasing morphological complexity. It seems that maturation of reprogrammed RGCs requires more than 3 weeks and possibly additional factors to ASCL1 in order to generate fully mature RGCs. This is perhaps not surprising given the much longer developmental time for macaques (gestation 183 days) compared with mice (gestation 21 days).

In order to be able to use the CMV promoter *in vivo* or in intact retinal explants, we need an alternative strategy to restrict vector cargo expression only to glia. MicroRNA-mediated silencing has been harnessed successfully across fields, such as cancer research^37^ and neuroscience^38^, to restrict vector genomes in targeted cells. By adding miR124 target sites at the 3’UTR of the vector cassette, we were able to de-target endogenous neurons from both primary dissociated cultures and retinal explants. In our experiments, 100% of the cells that expressed the mNG reporter in explants also expressed the MG marker Sox2. Importantly, and of great translational relevance, is that the addition of miR124 target sites did not affect the reprogramming yield of adult MG into RGCs. Overexpressing ASCL1 in MG from either adult or infant retina had the same effect in inducing neurogenic precursors and immature RGCs. Interestingly, the infant MG treated with CMV-ASCL1-mNG-miR124T had a greater yield in neurogenesis and seemed to transition faster from precursors to RGCs. We hypothesize that in the infant, ASCL1+ neurogenic precursors may upregulate miR-124, which in turn suppresses ASCL1 and pushes cells out of the precursor stage into neurons more effectively. This dynamic regulation of ASCL1 via miR-124 is reminiscent of ASCL1 oscillation in neural progenitors during development, where it is high in cycling progenitors but then drops as they exit the cell cycle and commit to neurons^39^.

From our studies across species, we conclude that cell cycle re-entry may not be a prerequisite for glia-derived neurogenesis. In the adult macaque retina, of all the ASCL1+ MG 5.3% re-entered the cell cycle (EdU+) and 5.4% may have directly transdifferentiated into HuC/D+ amacrine/RGC-like neurons (EdU-) within 3 weeks of incubation. In our previous *in vivo* studies, using both transgenic and vector-mediated ASCL1 overexpression in adult mice, we showed that only a subset of reprogrammed neurons originate from a proliferating precursor^9,12,24^. Although cell cycle re-entry may not be necessary for MG-mediated neurogenesis, and direct reprogramming from glia to neurons can be possible, it is still an important control to demonstrate EdU+ neurons of a given subtype that can be lineage traced to MG.

We believe this body of work offers significant insights that will inform subsequent studies on the regenerative potential of MG in the adult retina. Most importantly, we highlight the need for testing regenerative strategies on non-human primate or human tissues. The primary type of neurons generated by reprogramming macaque MG are immature RGCs, unlike mouse MG that generate bipolar cells when overexpressing ASCL1 both *in vitro* and *in vivo*^9–12^. It is interesting that MG from different species respond to reprogramming factors with different outcomes as to the cell types generated. This is not surprising however, since previous reports have identified several species-specific differences in retinal development^40^, which is likely influenced by distinct roles of bHLH transcription factors across species; for example ASCL1+ progenitors in mice produce all neuron classes except RGCs^41^. As such, using relevant models is critical for assessing the therapeutic potential of a regenerative strategy. Our data suggests that RGCs can be generated from an endogenous cell source, which we envision can benefit patients suffering from glaucoma and other optic neuropathies in the future.

## Acknowledgements

The authors would like to thank all the members of the Reh lab, especially our lab manager Catherine Ray for her extensive support, as well as the Bermingham-McDonogh lab for their valuable comments on the manuscript. We would also like to thank Christopher English for his support throughout tissue acquisition and distribution. All schematics were generated using Biorender.

## Author contributions

Conceptualization: MP, TR

Methodology: MP, JW, JH, LK, FK, IOL

Investigation & Visualization: MP, JW, JH, LK, FR, MBM, IOL, ARP, MPr

Funding acquisition: MP, TR

Project administration: MP, TR

Supervision: TR

Writing – original draft: MP, TR

Writing – review & editing: MP, JW, JH, LK, TR

## Materials and Methods

### Macaque retina harvesting

All retina samples were collected within 2 hours of necropsy at the Washington National Primate Research Center. Enucleated globes were dissected to remove the anterior chambers and where vitrectomy was challenging a digestion step with plasmin was implemented, followed by vitrectomy. The remaining eyecup was kept in oxygenated Ame’s medium and then transferred onto a dissection plate with PBS/glucose (12 mM). The retina was gently peeled off all underlying tissue and transferred into a container with fresh PBS/glucose for subsequent processing for either MACS or explant cultures.

### CD29-based magnetic activated cell sorting

The size of the retina varied across samples; therefore, retina was dissected into 5×5mm pieces, and 4-5 pieces were combined into a tube with 672μl PBS/glucose for dissociation with papain (0.2 mg/ml; Roche) for 30 min at 37°C without agitation. The tissue pieces were then washed 3 times with 500ul PBS/glucose and subsequently incubated with DNase I (200 U/ml in PBS/glucose) for 4 min at RT. PBS/glucose was removed and substituted with 400μl extracellular solution (ECS, 136 mM NaCl, 3 mM KCl, 10 mM HEPES, 11 mM glucose, 1 mM MgCl2 and 2 mM CaCl2, pH 7.4) before triturating the tissue using a firepolished glass pipette. 100μl biotinylated anti-CD29 human antibody (Miltenyi) was added to the cell suspension and incubated for 15mins at 4°C. 500μl ECS was added to each tube and centrifuged at 600xg for 10min at 4°C. The supernatant was discarded, and the pellet was resuspended in 200μl ECS using the firepolished glass pipette. The cell suspension was incubated with 50μl anti-Biotin magnetic beads (Miltenyi) for 15mins at 4°C, topped up with 700μl ECS and spun down at 600xg for 10min at 4°C. The supernatant was discarded, the pellet was resuspended in 1ml ECS and then passed through a cell strainer. The cell suspension was then added to a Large Cell Column (Miltenyi) previously rinsed with 500μl ECS. Once the cell suspension passes through the column by gravity flow each column was washed 3 times with 1ml ECS. Columns were then removed from the magnetic rack, loaded with 1ml ECS and pushed into a collection tube using the column plunger. To culture the CD29+ or CD29-fractions, cells were spun for 10mins at 400xg, resuspended in media (Neurobasal, 5%FBS, B27, N2, ITS, P/S, L-Glutamine, EdU) and plated on culture plates coated with matrigel.

### Retinal explant cultures

Retinas were dissected into 1×1cm pieces and placed on 6-well polycarbonate membrane transwells (Corning) with the inner limiting membrane facing upward. Retinal explants were cultured at 37°C in 5% CO_2_ for three weeks in DMEM/F-12 nutrient medium (ThermoFisher), supplemented with 0.1% BSA, 10 μM O-acetyl-l-carnitine hydrochloride, 1 mM fumaric acid, 0.5 mM galactose, 1 mM glucose, 0.5 mM glycine, 10 mM HEPES, 0.05 mM mannose, 13 mM sodium bicarbonate, 3 mM taurine, 0.1 mM putrescine dihydrochloride, 0.35 μM retinol, 0.3 μM retinyl acetate, 0.2 μM (±)-α-tocopherol, 0.5 mM ascorbic acid, 0.05 μM sodium selenite, 0.02 μM hydrocortisone, 0.02 μM progesterone, 1 μM insulin, 0.003 μM 3,3′,5′-triiodo-l-thyronine, 2,000 U penicillin, and 2 mg streptomycin (Sigma-Aldrich). The culture medium was changed every other day.

### Electrophysiology

Experiments were conducted on dissociated cells and whole mount preparations taken from dark-adapted retinal tissue of a fetal day 155 macaque that was made available through the Tissue Distribution Program of the National Primate Research Center at the University of Washington. All procedures were approved by the University of Washington Institutional Animal Care and Use Committee. Data were acquired at 10–kHz using a Multiclamp 700B amplifier (Molecular Devices), Bessel filtered at 3–kHz (900–CT, Frequency Devices), digitized using an ITC-18 analog-digital board (HEKA Instruments) and acquired using the Symphony acquisition software package (http://symphony-das.github.io). Freshly isolated retina was stored in oxygenated (95% O2/5% CO2) Ames medium (Sigma) at ∼32–34° C and, once under the microscope, tissue preparations were perfused by the same Ames solution at a rate of ∼8 mL/min. Isolated retinas were flattened onto poly-L-lysine slides (whole mount) as previously described (42, 43). Retinal neurons were visualized and targeted for cell-attached and/or whole-cell recordings using infrared light (>950 nm). All recordings used an intra-cellular solution containing (in mM): 123 K-aspartate, 10 KCl, 10 HEPES, 1 MgCl2, 1 CaCl2, 2 EGTA, 4 Mg-ATP and 0.5 Tris-GTP (∼280 mOsm; pH ∼7.2 with KOH). Biocytin hydrazide (1%; ThermoFisher) was added to the pipette solution to recover cellular morphology after recording, tissue fixation (4% PFA/PBS) and immunostaining (streptavidin Alexa Fluor 488; ThermoFisher).

### Lentiviral transduction

To transduce cell cultures, 2μl of the lentiviral prep was added in each well of a 24-well plate. After 48 hours the media was changed (Neurobasal, 1%FBS, B27, N2, ITS, P/S, L-Glutamine, EdU) and subsequently changed 50% every other day. To transduce retinal explants, 8-10μl of the lentiviral prep was pipetted directly on top of each explant and the explant media was supplemented with 1mM sodium butyrate.

**Table 2:** summary of lentiviral constructs and corresponding titer.

| lentiviral construct | TU/ml |
| --- | --- |
| LV/CMV>hASCL1[NM_004316.4]:IRES:TurboGFP | 3.99x10<sup>8</sup> |
| LV/CMV>EGFP:T2A:Puro-EF1a>mCherry | 4.49x10<sup>8</sup> |
| LV/CMV>hASCL1[NM_004316.4](ns):T2A:mNeon-Green:WPRE:miR124-3pT | 2.70x10<sup>8</sup> |
| LV/CMV>mNeonGreen:WPRE:miR124-3pT | 2.85x10<sup>8</sup> |

### Immunolabeling

Cells on coverslips were fixed with 4% PFA/PBS for 15mins and then kept in PBS at 4ºC. Retinal explants were fixed with 4% PFA/PBS for 30mins and then sequentially dehydrated with 10% sucrose/PBS for 1hour, 20% sucrose/PBS for 1hour, 30% sucrose/PBS overnight at 4ºC. Explants were then embedded in OCT compound before freezing. Frozen samples were sectioned at −20°C in 15-to 18-µM sections onto glass slides. Slides were then heated for 10 min on a slide warmer before staining or freezing at −20°C for long-term storage. For EdU detection the Click-iT™ EdU Imaging Kit (Thermo Fisher Cat # C10086) was used following manufacturer instructions. Primary antibodies were incubated overnight at 4ºC in blocking solution (5% normal horse serum, 5% bovine serum albumin, 1% TritonX-100 in PBS). The primary solution was removed, and slides were washed 3x with PBS for 10mins. Secondary antibodies were incubated in blocking solution for 90 min and slides were mounted using Fluoromount-G (SouthernBiotech).

**Table 3:** List of antibodies and corresponding dilution factor.

| Antibody | Source | Identifier | Dilution |
| --- | --- | --- | --- |
| chicken anti-GFP | Abcam | AB13970 | 1:1000 |
| mouse anti-HuC/D | Invitrogen | A-21271 | 1:200 |
| goat anti-OTX2 | R&D Systems | BAF1979 | 1:500 |
| rabbit anti-ASCL1 | Abcam | AB211327 | 1:300 |
| goat anti-SOX2 | Santa Cruz | SC-17320 | 1:200 |
| rabbit anti-RCVRN | Millipore | AB5585 | 1:500 |
| rabbit anti-SOX9 | Millipore | AB5535 | 1:500 |
| mouse anti-CRALBP | Abcam | AB15051 | 1:300 |
| mouse anti-GS | Millipore | MAB302 | 1:500 |
| rabbit anti-GFAP | Dako | z0334 | 1:500 |
| goat anti-BRN3 | Santa Cruz | sc-6026 | 1:200 |
| mouse anti-TUJ1 | Biolegend | 801202 | 1:500 |
| mouse anti-ISL1 | DSHB Hybridoma | 39.4D5 | 1:200 |
| mouse anti-mNG | Chromotek | 32f6-100 | 1:500 |
| rabbit anti-mNG | Chromotek | nfrb-100 | 1:500 |
| goat anti-ChAT | Millipore | AB144 | 1:200 |
| donkey anti-chick 488 | Jackson Immuno | 703-545-155 | 1:500 |
| donkey anti-goat 405 | Invitrogen | A48259 | 1:500 |
| donkey anti-goat 568 | Invitrogen | A11057 | 1:500 |
| donkey anti-goat 647 | Jackson Immuno | 705-605-147 | 1:500 |
| donkey anti-mouse 488 | Jackson Immuno | 715-605-150 | 1:500 |
| donkey anti-mouse 568 | Invitrogen | A10037 | 1:500 |
| donkey anti-mouse 647 | Jackson Immuno | 715-605-150 | 1:500 |
| donkey anti-rabbit 488 | Invitrogen | A21206 | 1:500 |
| donkey anti-rabbit 568 | Invitrogen | A100042 | 1:500 |
| donkey anti-rabbit 647 | Invitrogen | A31573 | 1:500 |

### Microscopy

Stained sections were imaged using a Zeiss LSM800 microscope. For quantification, images were taken with a 20x objective with at least 3 images taken per sample and condition. Images were analyzed and counted using FIJI. The colocalization of markers was verified by going through single-planes of the z-stack. Careful attention was paid to ensure there was a consistent pattern of marker overlap in each plane where the cell was present. Individual performing cell counting was blinded whenever possible.

### Single cell RNA sequencing

Cell cultures were processed for single-cell RNA sequencing by removing the media and washing twice with 0.04%BSA/DPBS. Cells were detached using 300μl Accutase with 20μl DNase I (10U/μl) at 37 ºC and then spun down at 500xg for 5min at 4ºC. Cell pellet was resuspended in 1 ml 0.04%BSA/DPBS and passed through cell strainer to remove any clumps. Library construction was performed using the Chromium Next GEM Single Cell 3′ version 3.1 (dual index) protocol and reagents according to the manufacturer’s instructions. Multiplexed libraries were sequenced using an Illumina NextSeq 500 using high-output 150 kits. Sequencing reads were demultiplexed and aligned to the *Macaca nemestrina* reference genome, Mnem_1.0, using Cellranger to obtain gene expression matrices for each sample. Gene expression matrices were read into R and analyzed with Seurat using a standard pipeline^42^. Briefly, cells were log-normalized, and the top 2000 highly variable features were used to perform principal component analysis. These components were used to identify clusters and embed cells in a 2D space using UMAP, and the expression of known marker genes was used to assign labels to each cluster.

### Cross Species Reference Mapping and Label Transfer

To compare our *Macaca nemestrina* dataset to the adult *Macaca fascicularis* and developing human fetal references, orthology tables were generated via Ensembl BioMart to identify a list of one-to-one orthologous genes conserved between each species. Datasets were subset to this list of conserved genes between each species and combined along with these features for further analysis. Seurat’s single-cell reference mapping was used to project our data onto the other reference’s space and perform label transfer. Transfer anchors were identified using FindTransferAnchors() with the first 35 principal components as the reference reduction, our data as the query, and either the human fetal or adult macaque atlas as the reference. These transfer anchors were then used to project cell type labels using TransferData() and embed cells in the reference UMAP using MapQuery().

### Pseudotime Analysis

After projecting our data onto the developing human fetal atlas, we used Monocle3 to construct pseudotime trajectories of the reprogrammed MG. Using the human fetal UMAP embedding, cells were ordered in pseudotime using order_cells() by setting the graph node that lied within the human fetal MG cluster as the starting node.

**Figure S1:**
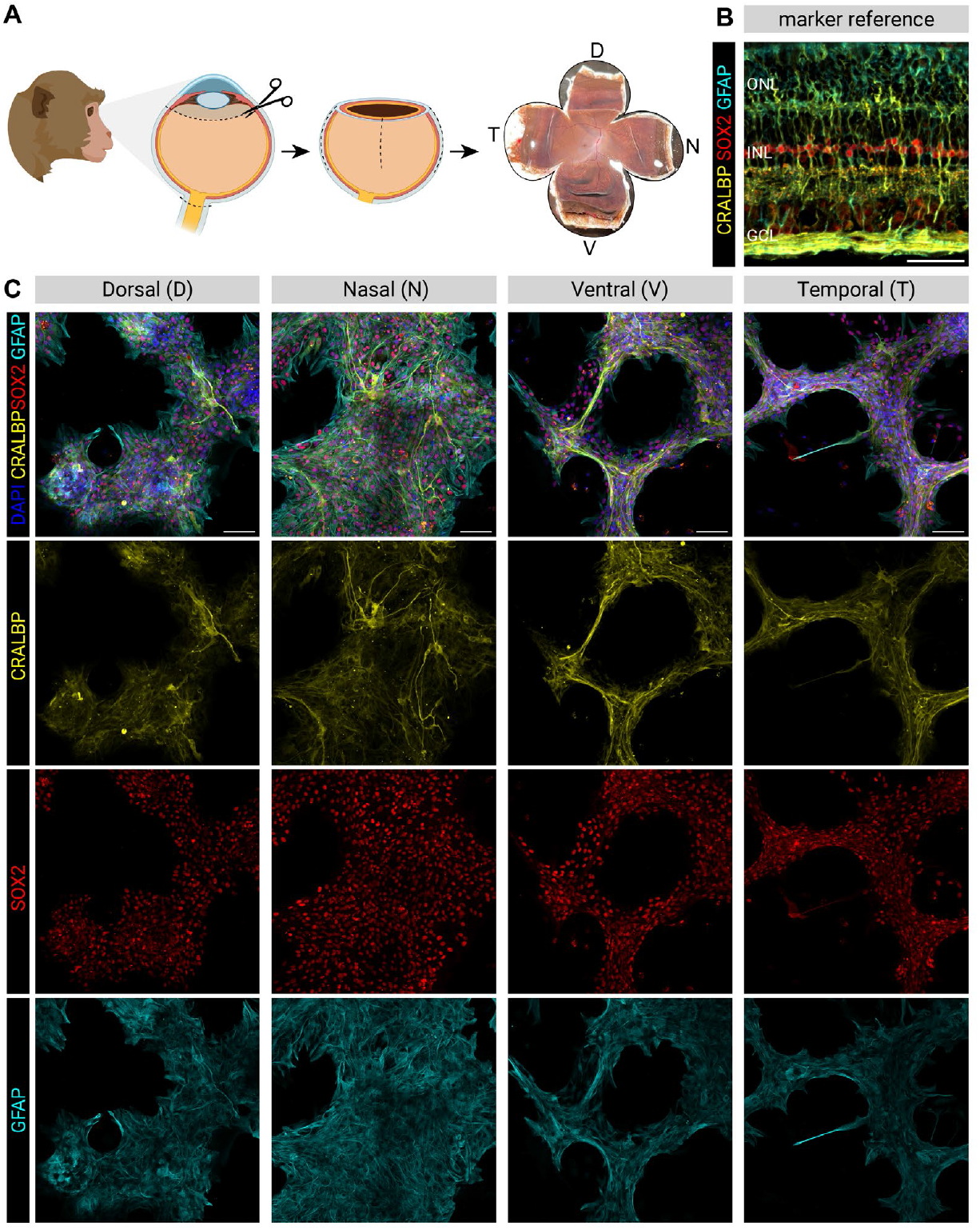
Müller glia can be cultured from all retinal regions of the macaque retina: (A) schematic diagram outlining the macaque eyecup dissection into dorsal [D], ventral [V], nasal [N] and temporal [T] quadrants, (B) fluorescence image of macaque retina cross section stained for glial markers, (C) fluorescence images of primary retinal cultures from each quadrant stained for glial markers as in (B). Scalebars: 50μm

**Figure S2:**
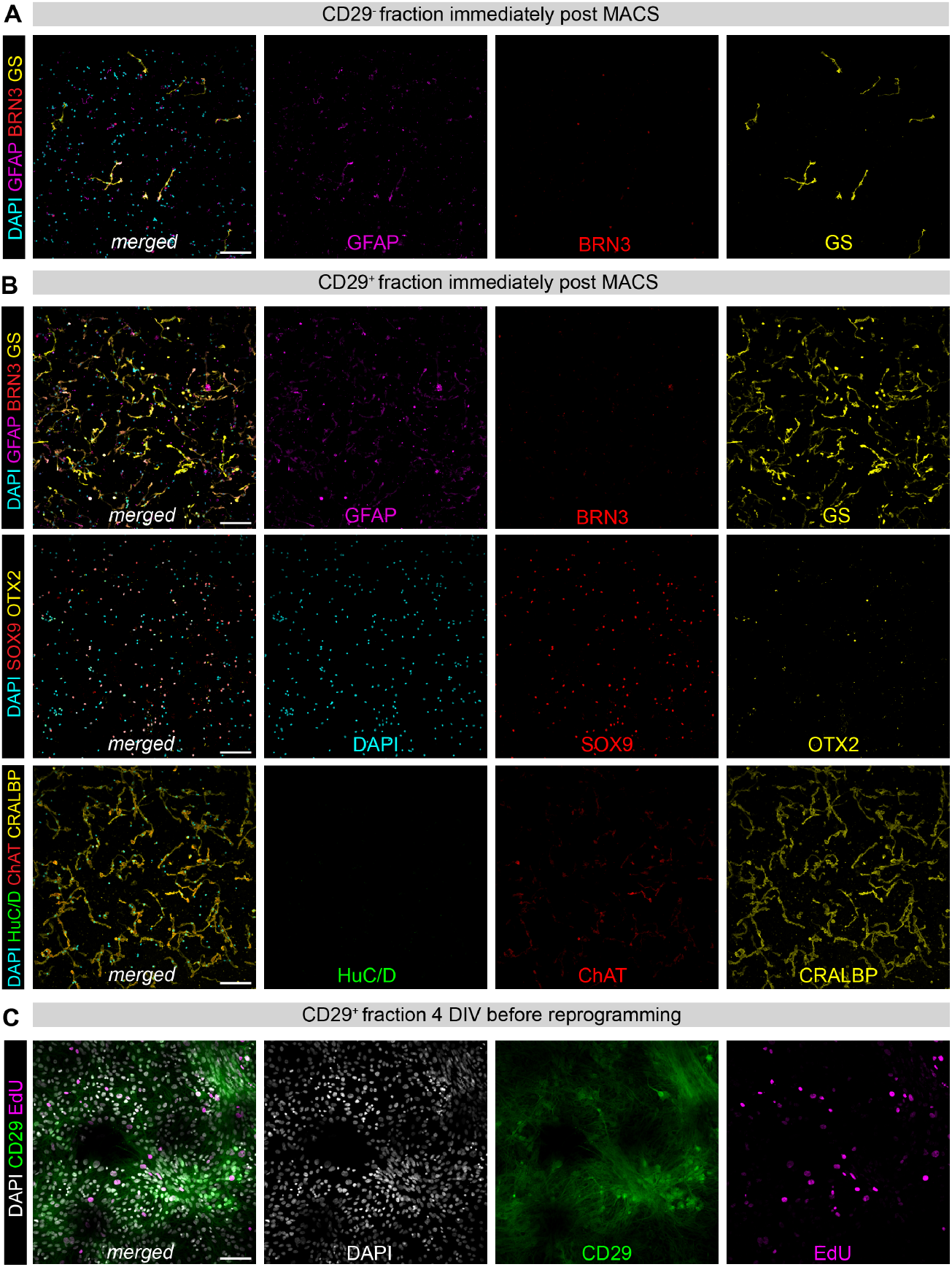
Marker characterization of cellular fractions post MACS: (A) fluorescence images of CD29-cellular fraction immediately post MACS stained for glial markers GS, GFAP and RGC marker BRN3, (B) fluorescence images of CD29+ cellular fraction immediately post MACS stained for glial and neuronal markers, (C) fluorescence images of CD29+ cellular fraction after 4 days *in vitro* (DIV) stained for CD29 antibody and proliferation marker EdU. Scalebars: 100μm

**Figure S3:**
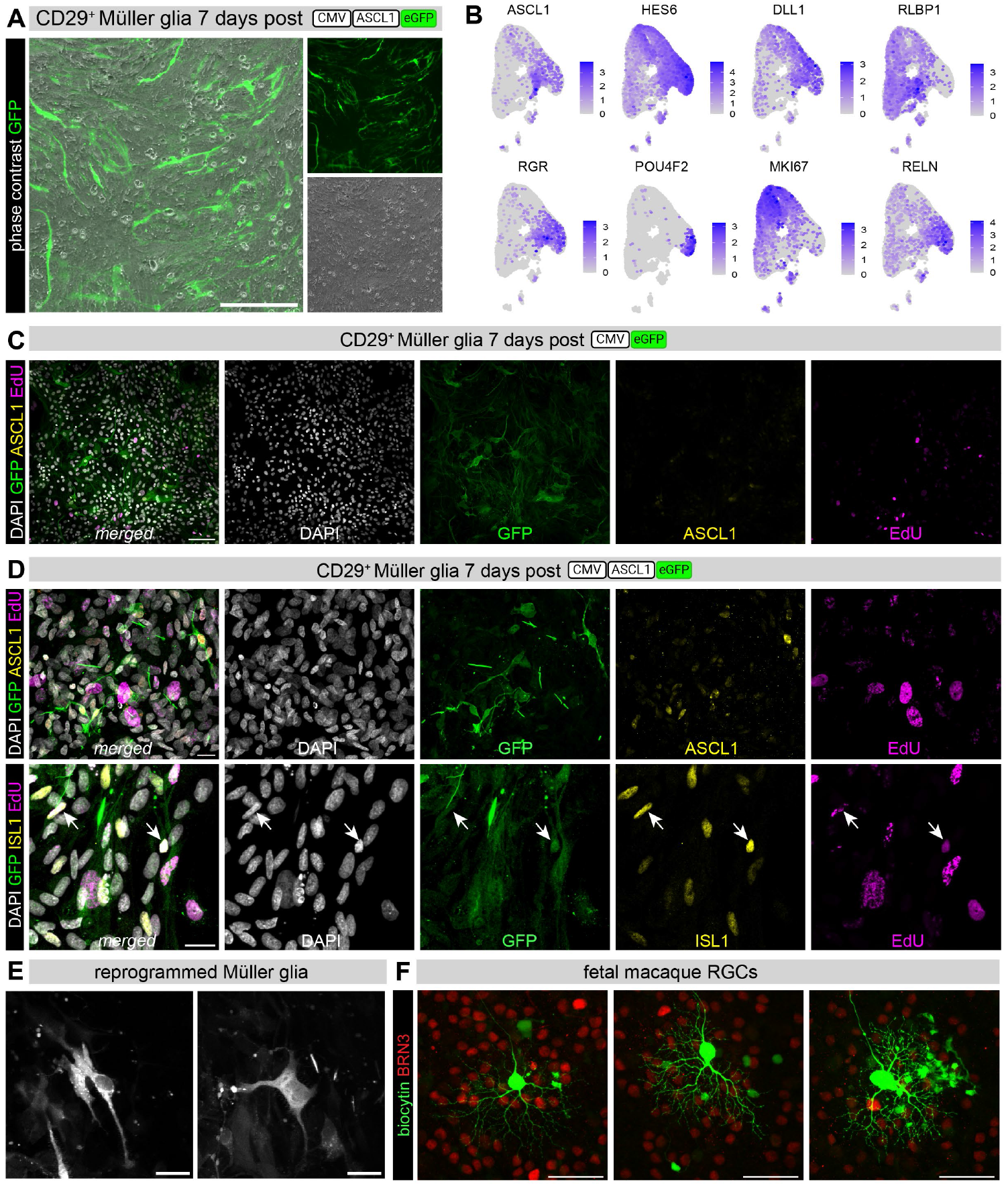
ASCL1 drives purified MG cultures to give rise to RGC-like neurons: (A) composite images of fluorescence and phase contrast of adult MG culture transduced with LV/CMV-ASCL1-eGFP, (B) feature plots of gene expression on integrated scRNA-seq data from adult MG treated with control CMV-eGFP and CMV-ASCL1-eGFP 7 days post transduction, (C) fluorescent images of adult MG culture treated with control CMV-eGFP labeled for GFP, ASCL1 and proliferation marker EdU, (D) fluorescent images of adult MG culture treated with control CMV-ASCL1-eGFP labeled for GFP, ASCL1, EdU and ISL1 with white arrows showing examples of GFP+ISL1+EdU+ cells 7 days post transduction, (E) fluorescent images of reprogrammed MG and (F) fetal macaque retina labeled with RGC marker BRN3 showing BRN3-RGCs previously patch-clamped and filled with biocytin. Scalebars:100μm for (A), 20μm for (C-E), 50μm for (E).

**Figure S4:**
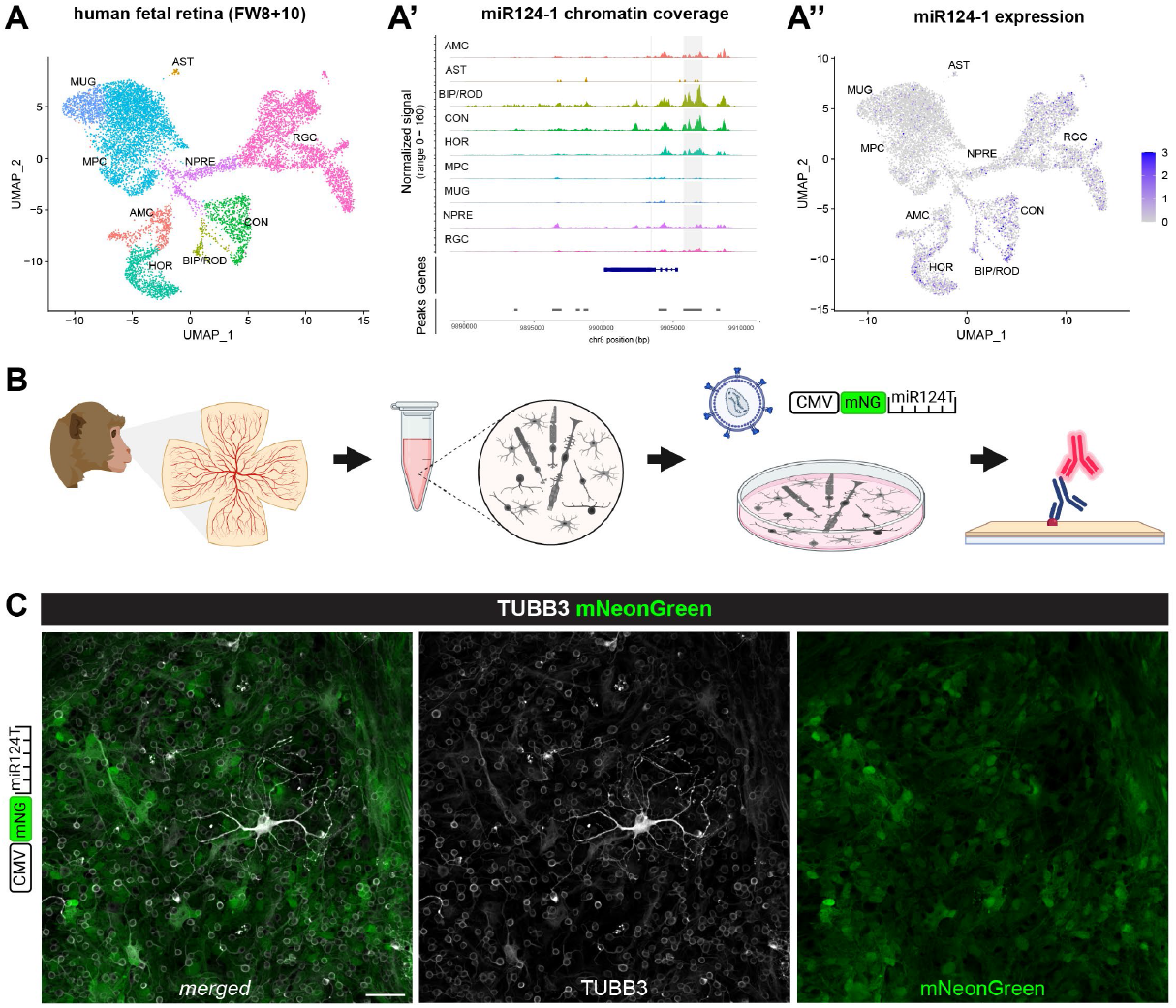
MicroRNA-mediated strategy to de-target endogenous neurons: (A) UMAP of Multiome sequencing data from human fetal retina at gestation week 8 and 10 with cell clusters identified according to common transcripts as previously published in ^15^, (A’) chromatin coverage plot for the microRNA-124-1 sequence of human fetal retina cell clusters highlighting the absence of open chromatin in glial cell clusters (AST:astrocytes, MUG:Muller glia) and MPC: multipotent progenitor cells, (A’’) feature plot of miR124-1 expression across human fetal retina from (A), (B) schematic diagram of experimental testing the LV/CMV-mNG-miR124T construct on a mixed primary culture from adult macaque retina, (C) fluorescence images of mixed primary culture transduced with LV/CMV-mNG-miR124T. Scalebar: 50μm

